# Zooplankton feeding behaviour and survival to toxic and non-toxic cyanobacteria during the seasonal bloom progression of a eutrophic lake

**DOI:** 10.1101/2025.10.08.681118

**Authors:** Pinelopi Ntetsika, Salomé Stauffer, Stefanie Eyring, Marta Reyes, Xuejian Wang, Elisabeth M.-L. Janssen, Francesco Pomati

**Author notes:** Corresponding author: Pinelopi Ntetsika.

## Abstract

Harmful cyanobacterial blooms pose increasing threats to aquatic ecosystems and human health; yet, the role of zooplankton grazing in regulating blooms remains understudied. We investigated the seasonal feeding behaviour and fitness consequences of feeding preferences in natural zooplankton communities for toxic (microcystin-producing) versus non-toxic cyanobacteria across temperature gradients in eutrophic Lake Greifen, Switzerland. We conducted monthly experiments from April to October 2023 to test the grazing behaviour of four zooplankton groups (daphnids, calanoid copepods, cyclopoid copepods, and microzooplankton) exposed to mixed diets of green algae and either toxic or non-toxic *Microcystis* strains at 15°C and 25°C.

Contrary to expectations of cyanobacteria avoidance, zooplankton exhibited predominantly non-selective grazing throughout the seasonal succession, consuming both toxic and non-toxic cyanobacteria at similar rates, regardless of temperature. Notably, during the peaks of phytoplankton abundance (April and September), mesozooplankton demonstrated a selective preference for cyanobacteria over green algae, particularly non-toxic strains. Temperature effects were subtle but revealed metabolic constraints: elevated temperatures occasionally triggered selective consumption of cyanobacteria in copepods, while fitness costs (survival) from exposure to toxic species were mostly restricted to transitional bloom periods and high-temperature conditions.

These findings suggest that toxic cyanobacteria may not always evade grazing pressure through secondary metabolite deterrent effects. Our results suggest that zooplankton communities can adapt and graze on cyanobacteria regardless of toxicity under the tested conditions, even during bloom conditions. These observations highlight the potential for zooplankton to interact with cyanobacterial populations, which may have implications for bloom prediction and management strategies, particularly under climate warming scenarios.

**Manuscript Highlights:**

- Zooplankton grazed on toxic and non-toxic cyanobacteria with similar effects across seasons in a eutrophic lake characterised by toxic blooms.
- Natural zooplankton communities showed no systematic avoidance of microcystin-producing cyanobacteria.
- High temperature effects on feeding selectivity were subtle and taxon-specific.
- Fitness costs from exposure to cyanobacteria (including microcystin-producing isolates) were rare and occurred only during transitional bloom periods.
- Results suggest that zooplankton communities may be adapted to cyanobacterial blooms, even when dominated by toxic species.

## 1. Introduction

Cyanobacterial blooms are a growing global concern due to their ecological, economic, and public health implications. Blooms degrade water quality, cause hypoxia, and release toxins that threaten both aquatic ecosystems and human health (Huisman et al., 2018; Paerl and Otten, 5 2013). The increasing frequency and intensity of cyanobacterial blooms are strongly associated with anthropogenic drivers, particularly nutrient enrichment and climate change (Glibert and Burford, 2017; Huisman et al., 2018; Visser et al., 2016), which promote conditions favourable for cyanobacteria — namely, high water temperature and high phosphorus and nitrogen levels during thermal stratification. As a result, current research has predominantly focused on these bottom-up abiotic drivers, which are more easily measured, manipulated in experiments, and controlled for environmental restoration. However, top-down biotic interactions, especially those involving zooplankton grazing, remain poorly characterised despite their potential role in shaping bloom dynamics (Ger et al., 2016; Ntetsika et al., 2025; Sweeney et al., 2022).

Zooplankton are expected to play a critical and complex role in modulating cyanobacterial bloom dynamics through their diverse feeding behaviours and physiological adaptations. Major zooplankton groups — cladocerans, cyclopoid and calanoid copepods, rotifers, and ciliates — exhibit distinct feeding strategies that affect their ability to consume and control cyanobacteria (Nandini and Sarma, 2023; Sommer et al., 2002). *Daphnia* species, although efficient generalist filter feeders, often experience reduced growth and increased mortality when exposed to toxic or filamentous cyanobacteria (Ger et al., 2016). In contrast, copepods can employ selective feeding mechanisms to avoid toxic prey (Schultz and Kiørboe, 2009), though recent evidence suggests reduced selectivity towards cyanobacteria under certain conditions (Novotny et al., 2023). Microzooplankton, including rotifers and ciliates, have demonstrated the capacity to graze on cyanobacteria and can be more effective in controlling blooms than larger zooplankton in some ecosystems (Bruno et al., 2012; Novotny et al., 6 2021; Sweeney et al., 2022).

The complexity of zooplankton-cyanobacteria interactions is further complicated by temperature, which can fundamentally alter both grazing behaviour and toxin sensitivity. Recent evidence shows that temperature modifies zooplankton responses to cyanobacteria through multiple pathways: temperature-dependent changes in metabolic demands can drive shifts toward more energy-rich herbivorous diets (Boersma et al., 2016), while simultaneously altering nutritional requirements in non-linear patterns across thermal gradients (Laspoumaderes et al., 8 2022; Ruiz et al., 2020). Critically, temperature appears to amplify toxicity, with multiple taxa showing enhanced sensitivity to cyanotoxins at elevated temperatures (Claska, 2002; Xiang et al., 2017), though these responses vary among zooplankton groups and toxin types (Nandini et al., 2019). This temperature-toxin synergy suggests that warming waters may exacerbate the challenges posed by toxic blooms to zooplankton communities.

Evolutionary response and phenotypic plasticity of zooplankton populations in their response to cyanobacteria add further complexity to understanding bloom dynamics. While toxic and morphologically defended cyanobacteria pose significant challenges to grazers, several zooplankton taxa can evolve or phenotypically adapt to tolerate toxins. Mechanisms of tolerance include reduced sensitivity to toxins, adjustments in feeding behaviour, and various avoidance strategies (Ger et al., 2016; Schaffner et al., 2019). Populations of *Daphnia* in eutrophic lakes with recurrent blooms have developed greater tolerance to cyanotoxins and can suppress toxic cyanobacteria more effectively (Ger et al., 2016; Nandini and Sarma, 2023). Copepods such as *Eudiaptomus gracilis* have demonstrated increased avoidance efficiency over time, suggesting potential for induced responses to toxic prey (Ger et al., 2011). These adaptations can occur rapidly — in the order of weeks — due to short generation times and high reproductive rates of many zooplankton species, such as ciliates and rotifers, which have shown tolerance to cyanotoxins and can act as effective grazers during peak bloom conditions (Davis et al., 2012; Dirren et al., 2017).

Despite this growing recognition of their potential to feed on toxic cyanobacteria, the interactive effects of temperature, toxicity, and adaptation on zooplankton-cyanobacteria interactions remain poorly understood. Few studies have disentangled which zooplankton taxa experience stronger selection pressure during cyanobacterial blooms and how this affects their grazing behaviour or potential to control blooms across thermal regimes. The methodological challenges of scaling laboratory findings to natural ecosystems, coupled with the taxonomic diversity and the complexity of multi-stressor interactions, have hindered a comprehensive understanding of their ecological importance.

In this study, we bridge this gap by experimentally crossing zooplankton functional diversity, seasonal adaptation dynamics, temperature-toxin interactions, and natural bloom progression in a eutrophic lake with frequent and toxic cyanobacterial blooms (Monchamp et al., 2016; Tellenbach et al., 2016; Wang et al., 2024) (**Fig. 1A-B**). We conducted monthly grazing assays from April to October using field-collected zooplankton communities — cladocerans, calanoid and cyclopoid copepods, and microzooplankton — from the eutrophic Lake Greifen in Switzerland. These grazers were exposed to mixed diets of cyanobacteria and green algae cultures, where cyanobacteria were either microcystin-producers or non-toxic strains, crossed with temperature treatments at 15°C and 25°C to assess behavioural and fitness effects (**Fig. 1C**).

**Figure 1:**
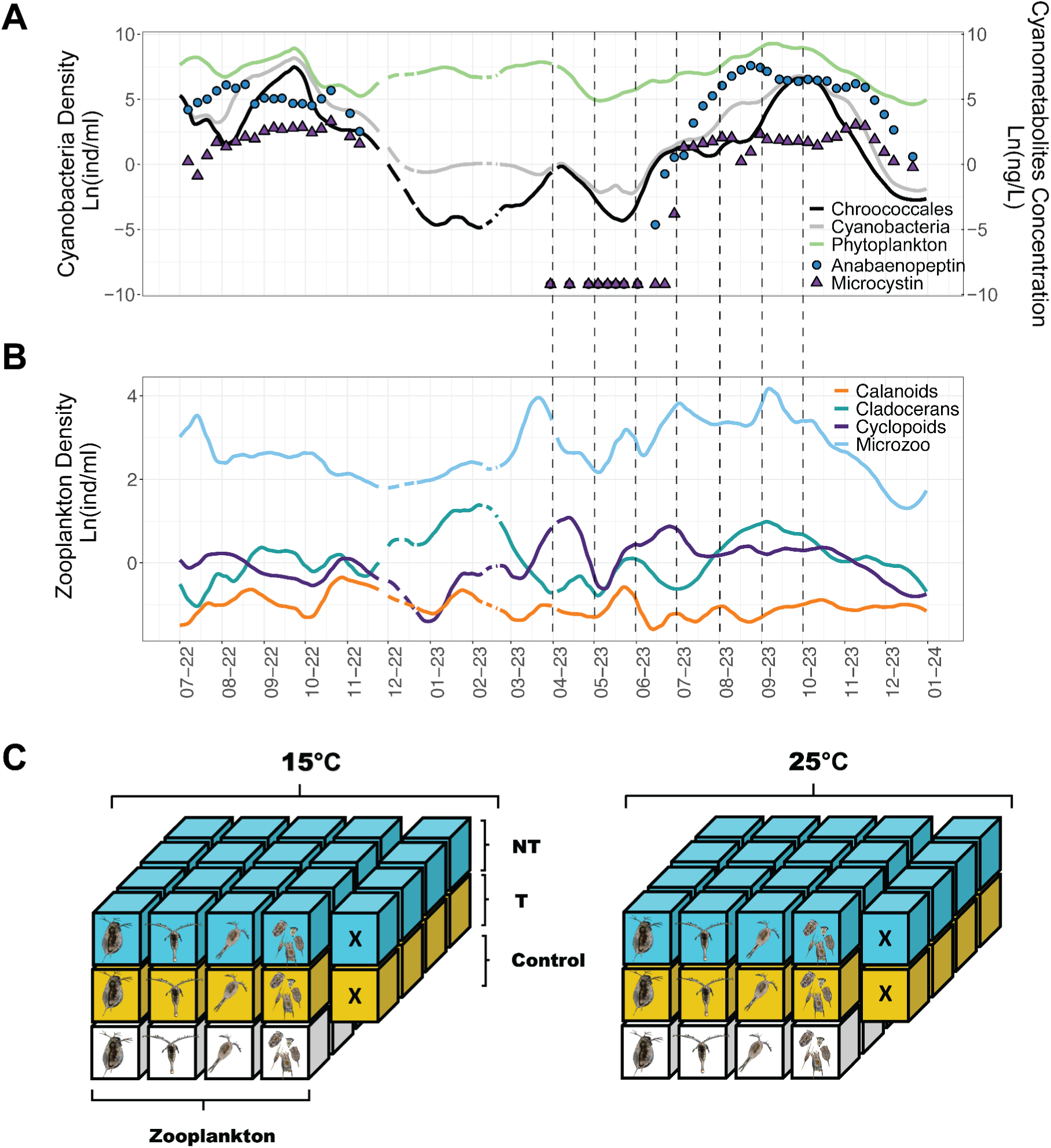
Temporal dynamics of plankton and cyanopeptides in Lake Greifensee, with an overview of the experimental design. **(A)** Time series of total phytoplankton, cyanobacterial and Chroococcales densities during the 2022-2023 growing season, alongside concentrations of the two major cyanobacterial metabolite classes: microcystins and anabaenopeptins. **(B)** Densities of the four target zooplankton groups—microzooplankton, daphnids (cladocerans), cyclopoid copepods, and calanoid copepods—across the same temporal window. Plankton data come from automated underwater microscopy (**Methods**); dashed lines indicate the timing of experimental tests. **(C)** Schematic of the experimental design, illustrating the factorial combination of food treatments (blue for non-toxic [NT], yellow for toxic [T], and white for green algae [Control]), temperature regimes (15°C and 25°C), and zooplankton species, including controls without grazers [X]. Note that with toxic, we mean microcystin-producing species (**Fig. S5**). Each treatment combination was replicated four times per month from April to October, totalling 112 experimental units.

Specifically, we ask: i) Which zooplankton taxa selectively graze or avoid cyanobacteria, and does toxin production make a difference? ii) How does increasing water temperature modify behavioural responses to toxic versus non-toxic cyanobacteria? iii) Does grazing behaviour shift across seasonal succession, and are zooplankton adapting to feed on cyanobacteria over time?

Based on a synthesis of the literature, we predict that feeding mechanisms will strongly influence behavioural responses and adaptive capacity relative to the presence of cyanobacteria (**Fig. 2**). Non-selective filter feeders should show greater sensitivity and higher fitness costs when exposed to cyanobacteria in their diet — especially toxic species — while selective grazers should incur fewer costs due to their discriminatory abilities. We expect increasing temperature to exacerbate toxic effects early in the season when zooplankton are naive to cyanobacteria, but adaptation should manifest as reduced fitness costs rather than altered feeding behaviour, with microzooplankton showing faster adaptation than larger grazers due to rapid turnover of generations.

**Figure 2:**
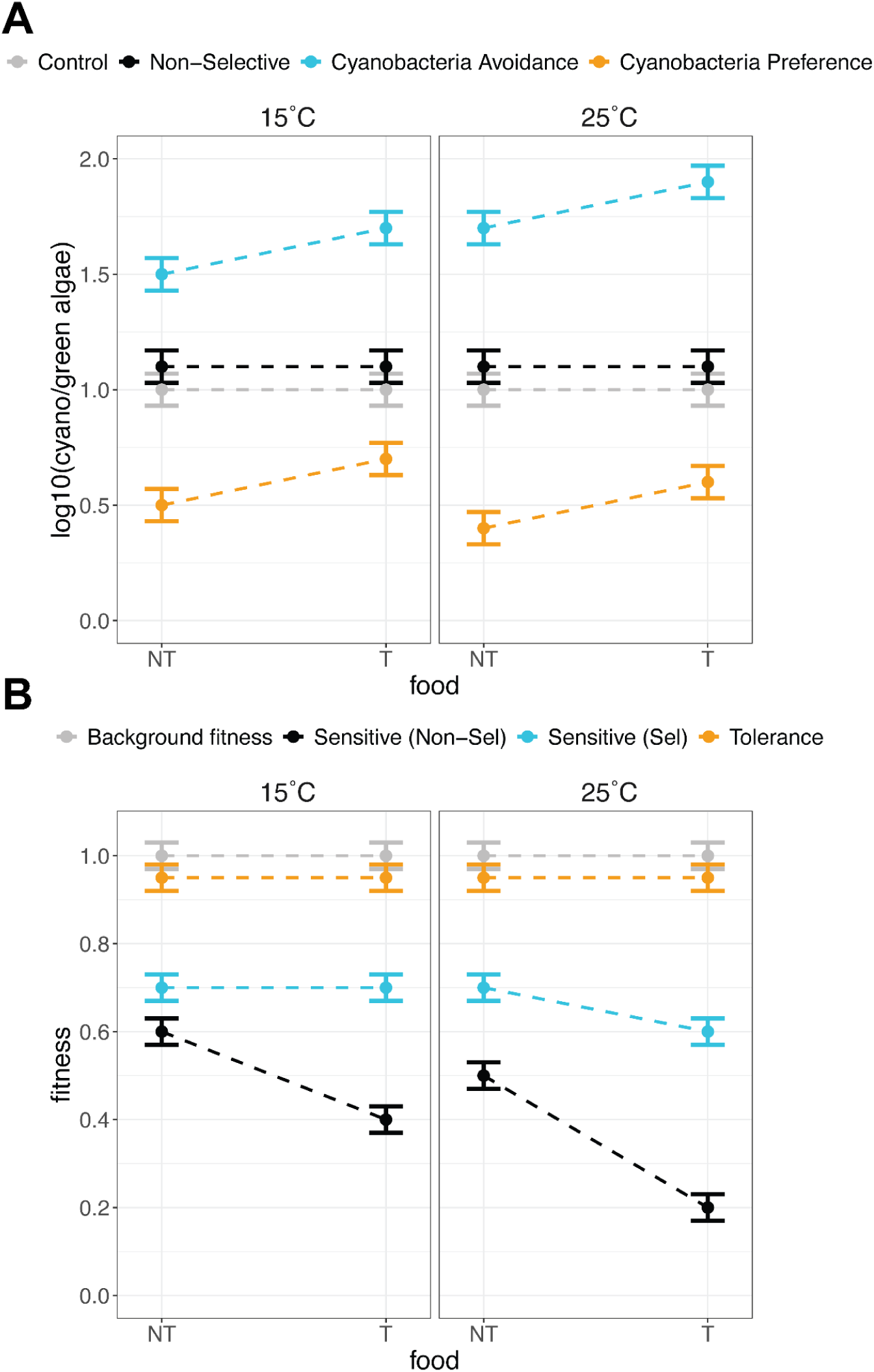
Expectations for zooplankton feeding selectivity and fitness under experimental treatments. Conceptual framework illustrating predicted changes in food selectivity and grazer fitness across food treatments and temperature conditions (**Methods**). Dots depict mean experimental values with their standard error. **(A) Feeding selectivity**. The Y-axis represents the Log(Response-Ratio)—food-ratio—of feeding preference: if the mean is significantly different from control levels, its position on the Y-axis indicates the type of behaviour. Non-selective grazers (in black) show no change in food-ratio relative to the control, regardless of treatment. Selective grazers deviate from the control depending on their feeding preference: a preference for cyanobacteria (in orange) decreases the food-ratio, while a preference for green algae (in blue) increases it. When zooplankton prefer cyanobacteria over green algae, the LogRR should be lower in the NT compared to the T treatment due to the presence of cyanotoxins. We expect the same pattern at both temperature regimes, with a slightly stronger selectivity (lower food-ratio) in the NT treatment. Avoidance of cyanobacteria (preference for green algae) should be stronger in the T treatment compared to the NT treatment. At high temperatures, we expect a greater preference for green algae (higher food-ratio) due to increased metabolic demands and the need for more nutritious food. **(B) Grazer fitness.** Non-selective grazers (in black) experience greater sensitivity to cyanobacteria and a fitness cost (measured as survival), with greater costs incurred when cyanobacteria are of the toxic type (T). Grazers that are sensitive to cyanobacteria but can selectively avoid them (in blue), maintain higher fitness levels compared to non-selective feeders. Increased metabolic demands from rising temperatures cause partial ingestion of low-quality (NT) or toxic (T) food and further decrease the fitness of both non-selective and cyanobacteria-avoidant grazers. Cyanobacteria-tolerant grazers (in orange) are expected to show higher fitness than the cyanobacteria-avoidant grazers, regardless of their feeding behaviour.

By spanning multiple functional groups across a full seasonal cycle with mixed-diet treatments and natural populations, our experimental design captures ecologically realistic feeding choices and complex preferences. We interpret our results in relation to seasonal plankton succession using high-frequency in situ data on zooplankton and cyanobacterial community composition, including concurrent measurements of cyanobacterial metabolites at the same sampling site. This approach provides critical evidence on how top-down control can modulate bloom trajectories, improving our mechanistic understanding of why blooms do not always expand even under favourable abiotic conditions — insight that is essential for predicting ecosystem response to continued warming and eutrophication.

## 2. Material & Methods

### 2.1 Study System

We conducted experimental trials using the natural zooplankton community of Lake Greifen, Switzerland (47.35 °N, 8.68 °E) from April to October 2023. Lake Greifen is a peri-alpine lake with a surface area of 8.45 km^2^ and a maximum depth of 32 m. The lake is monomictic and eutrophic with an annual average phosphorus load of 0.05 mg P/L (Eyring et al., 2025b).

We investigated the grazing response of four key zooplankton groups: daphnids, calanoid copepods, cyclopoid copepods, and microzooplankton. Monthly zooplankton samples were collected at either the lake’s deepest point (32 m) or our monitoring platform (maximum depth: 20 m) (Eyring et al., 2025b). Mesozooplankton (daphnids and copepods) were collected using a twin net (95 μm mesh size) and filtered through a 1 mm sieve (JBL Artemio 4), while microzooplankton were sampled with a 30 μm net at 10 m depth and sieved through a 150 μm sieve (JBL Artemio 4).

Lake Greifen’s plankton community shows a high turnover in composition across the seasonal succession and has a history of demonstrated cyanobacterial blooms (Monchamp et al., 2016a; Wang et al., 2024). During our 2023 sampling period, the phytoplankton community composition in spring was mostly dominated by small chlorophytes, cryptophytes, and dinoflagellates. In July, *Dinobryon* became the prominent genus, while from August to October, cyanobacteria prevailed, with primary representatives of the genus *Limnoraphis, Microcystis* or *Woronichinia*, among others. These observations come from a dual-magnification dark field underwater microscope installed in the lake (www.aquascope.ch) (Eyring et al., 2025b; Merz et al., 2021).

### 2.2 Experimental Design

#### 2.2.1 Zooplankton Target Groups

Zooplankton diversity remained relatively constant across the season, though notable shifts in relative abundance occurred (**Fig. S1**). The *Daphnia* genus in Lake Greifen primarily consists of *Daphnia pulicaria*, *D. cuculata*, *D. galeata*, *D. longispina* and their hybrids (Tellenbach et al., 2016) (**Table S1**). Calanoid copepods are predominantly represented by *Eudiaptomus gracilis,* while cyclopoid copepods include *Cyclops vicinus, C. abyssorum, C. bohater* and occasionally *Mesocyclops sp*. The microzooplankton community also exhibited high temporal variability. The main taxa present in the experiment included rotifers (*Polyarthra sp., Keratella quadrata, K. cochlearis*), ciliates (*Tinntinopsis sp., Coeleps sp., Tintinnidium sp.*) and nauplii (larval stages of copepods).

#### 2.2.2 Lake Data

To contextualise our hypotheses and experimental findings within the seasonal dynamics experienced by zooplankton, we compiled data on cyanobacterial metabolite profiles from Lake Greifen, alongside high-frequency records of zooplankton group densities, Chroococcales and total cyanobacteria from the lake’s monitoring station. We focused on two important classes of cyanobacterial metabolites: i) microcystins (MC-LR, MC-LA, MC-YR, MC-RR, [D-Asp3,(E)-Dhb7]MC-RR and [D-Asp3]MC-LR) and ii) anabaenopeptins (Anabaenopeptin A, Anabaenopeptin B and Oscillamide Y), which are linked with *Microcystis* (in Chroococcales group) and include the dominant metabolite compounds detected in Lake Greifen. For details on cyanobacterial metabolite analysis and data processing, refer to (Wang et al., 2024), while information regarding planktonic data acquisition and processing is provided in (Eyring et al., 2025b). Both zooplankton and cyanobacteria densities were calculated based on the taxonomic groups presented in **Table S1**. The Chroococcales group included mainly *Microcystis* sp.*, Aphanothece* sp.*, Aphanocapsa* sp. Data on plankton density were collected at an hourly resolution using automated underwater microscopy and averaged to daily values (Eyring et al., 2025c). Metabolite concentrations were measured weekly from April to November 2022 and from March to December 2023 (Wang et al., 2024).

#### 2.2.3 Treatments

Zooplankton were exposed to three diet treatments: (i) toxic (T), (ii) non-toxic (NT), and (iii) a control (CTRL) (**Fig. 1C**). The T and NT treatments consisted of a 1:1 mixture of green algae and cyanobacteria based on carbon content, while the control contained only green algae. Carbon content per L of culture was estimated by calculating cell biovolume from cell measurements and converting it to carbon using published conversion factors (Hillebrand et al., 1999; Li et al., 2022; Rocha and Duncan, 1985; Schlesinger et al., 1981; Schlesinger and Shuter, 1981). Culture carbon concentrations were then calibrated against fluorescence (RFU) measurements to standardise diets to equal carbon contributions. Final diet concentrations were adjusted to 200 RFU, corresponding to approximately 9.18 mg C/L, based on pilot studies.

All treatments included two green algae species: *Scenedesmus acuminatus* (strain SAG 38.81), a crescent-shaped unicellular alga (mean cell length: 16.7 μm; width: 3.6 μm), and *Chlorella vulgaris* (strain EPSAG 211-11b), a round unicellular alga (cell diameter: 1.7-4.3 μm). Both species are considered high-quality food sources due to their nutritional value and easily digestible morphology (Nandini and Sarma, 2023).

The toxic treatment included two microcystin-producing *Microcystis aeruginosa* strains: PCC 7806 and UV006. The non-toxic treatment contained two microcystin-deficient strains: a Microcystis strain isolated from Lake Greifen (∼10 years ago) and Microcystis B-(McyB knock-out). All *Microcystis* cultures consisted of single, round cells (mean diameter: 3.8 μm, range: 2.3-5.2 μm), which, due to their non-colonial form, should not have imposed morphological constraints on the grazers.

All diet treatments had a final concentration of 9.18 mg C/L, aiming for a 1:1 cyanobacteria-to-green algae carbon content ratio (except for the control, which contained only green algae). While initial deviations from the target ratio occurred across months, within each monthly trial, the variation in concentration was comparable (**Fig. S2**). A control treatment, including only the phytoplankton taxa (T and NT) in the absence of grazers, was used to account for changes in food concentration due to algal competition. Each diet treatment was tested under two temperature conditions (15°C and 25°C) to assess the impact of increasing zooplankton metabolic rates on food consumption and feeding behaviour. Each treatment was replicated four times, resulting in 112 experimental units (**Fig. 1C**). The experiment was conducted monthly from April to October to examine potential physiological or behavioural adaptations in grazer responses to cyanobacteria over the seasonal progression.

#### 2.2.4 Workflow

Lake water was collected from the surface and filtered through a glass-fibre filter (GFF, Sartorius), followed by a particle retention filter (0.45-μm, Sartobran) to obtain particle-free water for the experiment. Microzooplankton density was standardised to 405 ± 327 individuals per 100 mL of filtered lake water based on the typical summer densities in Lake Greifen (Merz et al., 2023), though actual densities varied across months due to a calculation error (see **Fig. S3**).

We acclimated all zooplankton groups overnight at 20°C. Microzooplankton were placed in the dark to graze on residual phytoplankton that could not be manually separated. After 22 hours of acclimation, calanoids, cyclopoids, and daphnids were manually sorted from mesozooplankton samples under a stereomicroscope and rinsed with tap water to remove any residual phytoplankton. Five individuals of each group were then transferred to 100 mL of lake-filtered water. Pre-tests confirmed that this concentration yielded a measurable grazing signal (decrease in relative fluorescence units, RFU) within 24 hours. Only phenotypically healthy individuals were selected, allowing for variation in size and reproductive stage.

Initial concentrations of green algae and cyanobacteria were measured using a plate reader (Cytation5, BIOTEK High Throughput Imager) at the start of the experiment (Measurement 1). Concentrations of green algae were approximated using chlorophyll a (chl-a) fluorescence (excitation: ∼586 nm, emission: ∼647 nm; reported in RFU), while cyanobacterial concentrations were approximated using phycocyanin fluorescence (excitation: ∼445 nm, emission: ∼685 nm; reported in RFU). Experimental jars were randomly placed in two temperature-controlled chambers (15°C/25°C), which were alternated monthly to avoid chamber effects. All jars were kept in the dark during incubation to limit algal growth and nutrient recycling (Ger et al., 2019). Subsequent fluorescence measurements of chl-a and phycocyanin were taken after 16, 19, and 23 hours (measurements 2-4) with thorough mixing before each reading to ensure homogeneity.

After 23 hours, zooplankton fitness was assessed as survival by counting alive and dead individuals, including potential offspring (only observed in *Daphnia*). Microzooplankton fitness was quantified by fixing a 10 mL subsample with Lugol’s iodine solution and counting phenotypically intact individuals under a microscope.

### 2.3 Data Analysis

#### 2.3.1 Selective Grazing

To evaluate feeding preferences toward specific prey, we analysed changes in the relative concentration of cyanobacteria and green algae. For that, we used the log_10_-transformed ratio of phycocyanin to chl-a RFU after 19 hours (the most appropriate time period after inspecting the temporal trends in the experiment) - we call this log_10_-ratio “food-ratio”. A higher food-ratio (indicating more phycocyanin) reflects a greater relative abundance of cyanobacteria, while a lower food-ratio (indicating more chl-a) reflects a greater relative abundance of green algae (**Fig. 2**). We used ordinary least squares regression (described below) to model the food-ratio as a function of food treatment (2 levels: T, NT), temperature (2 levels: 15°C, 25°C), zooplankton group (5 levels: 4 grazer groups + control), and their interaction using multiple linear models. In addition, we corrected for differences in the initial food density and survival by incorporating two covariates: i) the cyanobacteria concentration (phycocyanin RFU) at the beginning of the experiment and ii) the number of alive individuals at the end of the experiment (scaled with min-max normalisation). To address our primary goal of understanding how food selectivity changes across the seasonal progression, we separately analysed each experimental trial (month), recognising the potential physiological or genetic differences in the zooplankton community across time. We constructed 7 distinct models, corresponding to the trials conducted from April to October.

#### 2.3.2 Absence of Grazing

An absence of change in the food-ratio can occur under two distinct scenarios: (1) both food types (chl-a and phycocyanin) increase proportionally, which may reflect an absence of grazing, or (2) decrease proportionally, if non-selective grazing removes both food types at similar rates. To distinguish between these possibilities and avoid misinterpretation of the selective grazing analysis, we examined the absolute changes in each pigment separately (**Fig. S4**). Our objective was to identify cases where the lack of change in the food ratio was due to a genuine absence of grazing activity. To achieve this, we calculated the net change in chl-a and phycocyanin concentrations from the beginning of the experiment to the end of the 19-hour incubation period. We refer to this difference as the *net grazing effect* ([RFU_19_−RFU_0_]). To test whether the mean net grazing effect differed significantly between grazing treatments and the control, we fitted a linear model for each pigment. The model included the fixed effects of food treatment (2 levels: T, NT), temperature (2 levels: 15°C, 25°C), and zooplankton group (5 levels: 4 grazer groups + control). As with the selective grazing analysis, the data were analysed separately for each experimental month. A statistically significant increase in both chl-a and phycocyanin under a zooplankton treatment was interpreted as evidence for the absence of grazing in that treatment (**Fig. S4**).

#### 2.3.3 Fitness

Zooplankton fitness was assessed via survival, measured as the number of alive individuals at the end of the experiment. Survival was modelled as a function of the zooplankton group (4 levels: grazer groups), food treatment (3 levels: T, NT and control), temperature (2 levels: 15°C, 25°C) and their interaction using a linear model. As with the selective grazing analysis, separate models were constructed for each experimental trial (April-October). Due to large differences in their initial densities (**Fig. S3**) and replication rates, mesozooplankton and microzooplankton densities were min-max normalised before analysis.

##### Statistical Testing and Post-Hoc Analysis

To determine which treatments had significant effects, we performed type III analysis of variance (ANOVA) on the fitted models to test the overall significance of each treatment and their interactions. When significant effects were detected, we conducted pairwise post hoc comparisons. These pairwise comparisons were adjusted for multiple testing using Tukey’s or Dunnet’s method to control the family-wise error rate. All statistical analyses were performed in R v4.4.1, using the stats (v4.5.1) and emmeans (v1.11.1) packages.

## 3. Results

### 3.1 Plankton and Secondary Metabolites

Our seasonal sampling campaign effectively captured the fluctuations in plankton community dynamics and cyanopeptide concentrations throughout 2023. Chroococcales and total cyanobacteria in Lake Greifen exhibited two periods of interest during the ecological succession: the first in April, coinciding with our initial experimental trial, with low cyanobacteria densities (despite a small peak of Chroococcales), and the second in October, corresponding to our final trial (**Fig. 1A**). Cyanobacterial densities were lowest between January-March and again during May-June, before steadily increasing through the summer and autumn months.

The temporal dynamics of secondary metabolites closely tracked cyanobacterial abundance patterns but with notable delays. Concentrations of both microcystins and anabaenopeptins remained low between April and June, then increased substantially during the second phase of the bloom. Anabaenopeptins peaked in August while microcystins reached maximum concentrations in November, with both metabolite classes remaining elevated from July through December (**Fig. 1A**).

Zooplankton community succession showed distinct group-specific patterns (**Fig. 1B**). Microzooplankton densities peaked before our April trial and again between July-October, with lowest abundances in May and June. Daphnid densities remained relatively low from April through July before peaking in September. Cyclopoid copepods showed the highest densities in April, declined in May, then maintained elevated numbers through October. Calanoid copepods consistently exhibited the lowest densities among all zooplankton groups, remaining relatively sparse from April through October with only a modest peak in May.

### 3.2 Feeding Behaviour and Grazing Selectivity

Across most months and experimental conditions, zooplankton exhibited non-selective grazing behaviour, maintaining food-ratios similar to control treatments regardless of temperature or food toxicity (**Table S2**, **Fig. 3**). This pattern held consistently for all zooplankton groups, indicating that neither food treatment (toxic vs. non-toxic cyanobacteria) nor temperature (15°C vs. 25°C) systematically influenced feeding preferences. However, two months revealed notable deviations from this general pattern: April and September (**Table S2, Fig. 3**).

**Figure 3:**
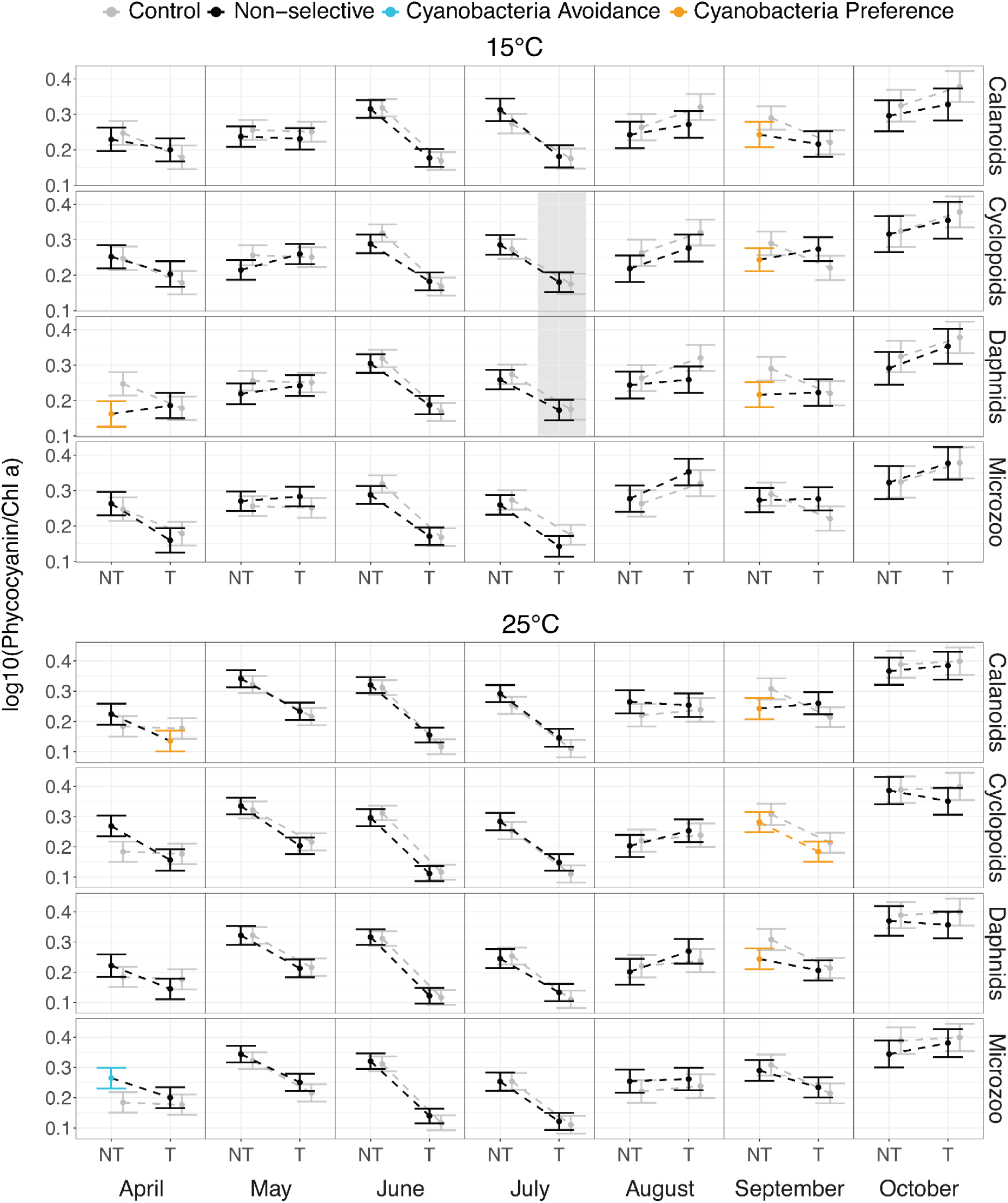
Predicted food-ratio (cyanobacteria/green algae) based on grazing selectivity analysis. Each panel shows a zooplankton group (rows) over time (columns), with linear-model predictions shown for each combination of food type and temperature. **Grey dots** indicate the mean of control levels (no grazers), while **colored dots** represent the mean treatments with zooplankton grazing under different food conditions: non-toxic (NT) and toxic (T) cyanobacteria. The color of the dots corresponds to the expected food-ratio outcomes under different zooplankton grazing behaviors (**see** Fig. 2 **A**); specifically blue and orange colors denote significant three-way interactions (*zooplankton × food × temperature*) identified through post-hoc testing. **Grey boxes** indicate cases of no grazing (see methods and **Fig. S4**). Error bars indicate 95% confidence intervals of predicted means. All statistically significant pairwise comparisons are detailed in **Table 1**.

**Table 1:** Results of significant post-hoc tests for the selective grazing analysis. Comparisons are organised by month (separate models) and treatment interaction. P-values are reported for each significant interaction, with values below 0.05 considered statistically significant.

| Contrast | Food | Temperature | Estimate | SE | Df | t.ratio | p.value |
| --- | --- | --- | --- | --- | --- | --- | --- |
| April |  |  |  |  |  |  |  |
| *Zoo* |  |  |  |  |  |  |  |
| Daphnids - Control |  |  | -0.037 | 0.017 | 58 | -2.207 | 0.031 |
| *Zoo:Food* |  |  |  |  |  |  |  |
| Daphnids - Control | NT |  | -0.045 | 0.021 | 58 | -2.108 | 0.039 |
| *Zoo:Temperature* |  |  |  |  |  |  |  |
| Daphnids - Control |  | 15 | -0.056 | 0.020 | 58 | -2.786 | 0.007 |
| *Zoo:Food:Temperature* |  |  |  |  |  |  |  |
| Daphnids - Control | NT | 15 | -0.102 | 0.026 | 58 | -3.907 | <0.001 |
| Microzoo - Control | NT | 25 | 0.057 | 0.028 | 58 | 2.083 | 0.042 |
| Calanoids - Control | T | 25 | -0.097 | 0.042 | 58 | -2.296 | 0.025 |
| May |  |  |  |  |  |  |  |
| *Temperature* |  |  |  |  |  |  |  |
| 15 - 25 |  |  | -0.031 | 0.006 | 58 | -5.150 | <0.001 |
| June |  |  |  |  |  |  |  |
| *Food* |  |  |  |  |  |  |  |
| NT - T |  |  | 0.157 | 0.006 | 58 | 27.307 | <0.001 |
| July |  |  |  |  |  |  |  |
| *Food* |  |  |  |  |  |  |  |
| NT - T |  |  | 0.111 | 0.012 | 57 | 9.177 | <0.001 |
| September |  |  |  |  |  |  |  |
| *Zoo* |  |  |  |  |  |  |  |
| Calanoids - Control |  |  | -0.055 | 0.023 | 58 | -2.364 | 0.021 |
| Cyclopoids - Control |  |  | -0.062 | 0.027 | 58 | -2.298 | 0.025 |
| Daphnids - Control |  |  | -0.048 | 0.014 | 58 | -3.393 | 0.001 |
| ***Zoo:Food*** |  |  |  |  |  |  |  |
| Calanoids - Control | NT |  | -0.098 | 0.027 | 58 | -3.561 | <0.001 |
| Cyclopoids - Control | NT |  | -0.087 | 0.030 | 58 | -2.938 | 0.005 |
| Daphnids - Control | NT |  | -0.076 | 0.018 | 58 | -4.189 | <0.001 |
| ***Zoo:Food:Temperature*** |  |  |  |  |  |  |  |
| Calanoids - Control | NT | 15 | -0.089 | 0.031 | 58 | -2.832 | 0.006 |
| Cyclopoids - Control | NT | 15 | -0.102 | 0.035 | 58 | -2.910 | 0.005 |
| Daphnids - Control | NT | 15 | -0.085 | 0.024 | 58 | -3.484 | <0.001 |
| Calanoids - Control | NT | 25 | -0.107 | 0.033 | 58 | -3.260 | 0.002 |
| Cyclopoids - Control | NT | 25 | -0.073 | 0.033 | 58 | -2.198 | 0.032 |
| Daphnids - Control | NT | 25 | -0.067 | 0.025 | 58 | -2.697 | 0.009 |
| Cyclopoids - Control | T | 25 | -0.080 | 0.034 | 58 | -2.347 | 0.022 |
| October |  |  |  |  |  |  |  |
| ***Temperature*** |  |  |  |  |  |  |  |
| 15 - 25 |  |  | -0.036 | 0.010 | 56 | -3.518 | <0.001 |

During April, when cyanobacteria were at their spring peak but toxins were absent from the lake (**Fig. 1A**), three zooplankton groups displayed selective feeding behaviours. Daphnids significantly reduced their food ratio in the non-toxic treatment at 15°C (mean change = -0.1; SE = 0.026; p < 0.001), indicating a clear preference for cyanobacteria over green algae (**Table 1**, **Fig. 3**). Calanoid copepods showed a similar cyanobacteria preference, but specifically in the toxic treatment at 15°C rather than the non-toxic treatment (mean change = -0.09; SE = 0.04; p = 0.025) (**Table 1**, **Fig. 3**). Conversely, microzooplankton—a heterogeneous group undergoing species turnover early in the season with few ciliates present (**Fig. S1**)—exhibited the opposite response, increasing their food ratio in the non-toxic treatment at 25°C (mean change = 0.057; SE = 0.028; p = 0.042) and thus preferring green algae over cyanobacteria (**Table 1**, **Fig. 3**).

September presented the most pronounced selective feeding responses across the zooplankton community (**Fig. 3**). During this period, when cyanobacteria dominated the phytoplankton community and toxin levels peaked (**Fig. 1A**), all mesozooplankton groups demonstrated significant preferences for cyanobacteria over green algae, but primarily when the cyanobacteria were non-toxic. Daphnids, calanoid copepods, and cyclopoid copepods all exhibited significantly lower food ratios in the non-toxic treatment at both temperatures, indicating systematic selective grazing against green algae and toward cyanobacteria (**Table 1**, **Fig. 3**). Additionally, cyclopoid copepods showed similar selective behaviour in the toxic treatment, but only at 25°C, suggesting that elevated temperature may influence their willingness to consume toxic prey (**Table 1**, **Fig. 3**).

An important exception to typical feeding patterns occurred in July, when both cyanobacteria abundance and toxin concentrations were rapidly increasing (**Fig. 1A**). Under these conditions, we observed complete feeding suppression rather than selective grazing. Specifically, both cyclopoid copepods and daphnids ceased feeding entirely when presented with toxic cyanobacteria at 15°C, as evidenced by significant increases in both chlorophyll a and phycocyanin concentrations relative to controls (**Fig. S4**). This feeding suppression occurred exclusively under the combination of toxic food and lower temperature, suggesting that environmental toxicity can completely inhibit grazing behaviour under certain conditions.

Temperature effects on feeding behaviour were generally subtle but revealed interesting patterns, with copepods at 25°C showing grazing selectivity towards cyanobacteria in April and September (**Table 1**, **Fig. 3**). This pattern suggests that higher temperatures may increase metabolic demands sufficiently to override typical feeding preferences, leading zooplankton to consume available food regardless of potential toxicity.

### 3.3 Survival and Fitness Responses

Zooplankton survival remained remarkably consistent across most experimental conditions, with no systematic differences between toxic and non-toxic food treatments or between the two temperature regimes tested (**Table 2**, **Table S3, Fig. 4**). Two primary exceptions to this pattern occurred in June and July. These months mark a transitional period when toxins are beginning to accumulate in the lake and cyanobacterial densities recover from their seasonal minimum (**Fig. 1A**). In June, both copepod groups experienced reduced survival, but in opposite treatments. Calanoid copepods showed significantly lower survival when fed toxic cyanobacteria at 25°C (mean scaled difference = 1.484; SE = 0.599; p = 0.041), while cyclopoid copepods exhibited reduced survival in the non-toxic treatment at the same temperature (mean scaled difference = 1.533; SE = 0.625; p = 0.043) (**Table 2**, **Fig. 4**). The restriction of these fitness costs to the elevated temperature treatment suggests that thermal stress may have exacerbated the effects of bioactive compounds (the average lake water temperature was 17°C in that period), whether the known microcystins in the toxic treatment or potentially unknown secondary metabolites produced by the non-toxic strains (**Fig. S5**). In July, calanoid copepods exhibited significantly reduced survival when fed toxic cyanobacteria, with the temperature effect not being significant, while all other zooplankton groups maintained baseline survival rates regardless of treatment (**Table 2**).

**Figure 4:**
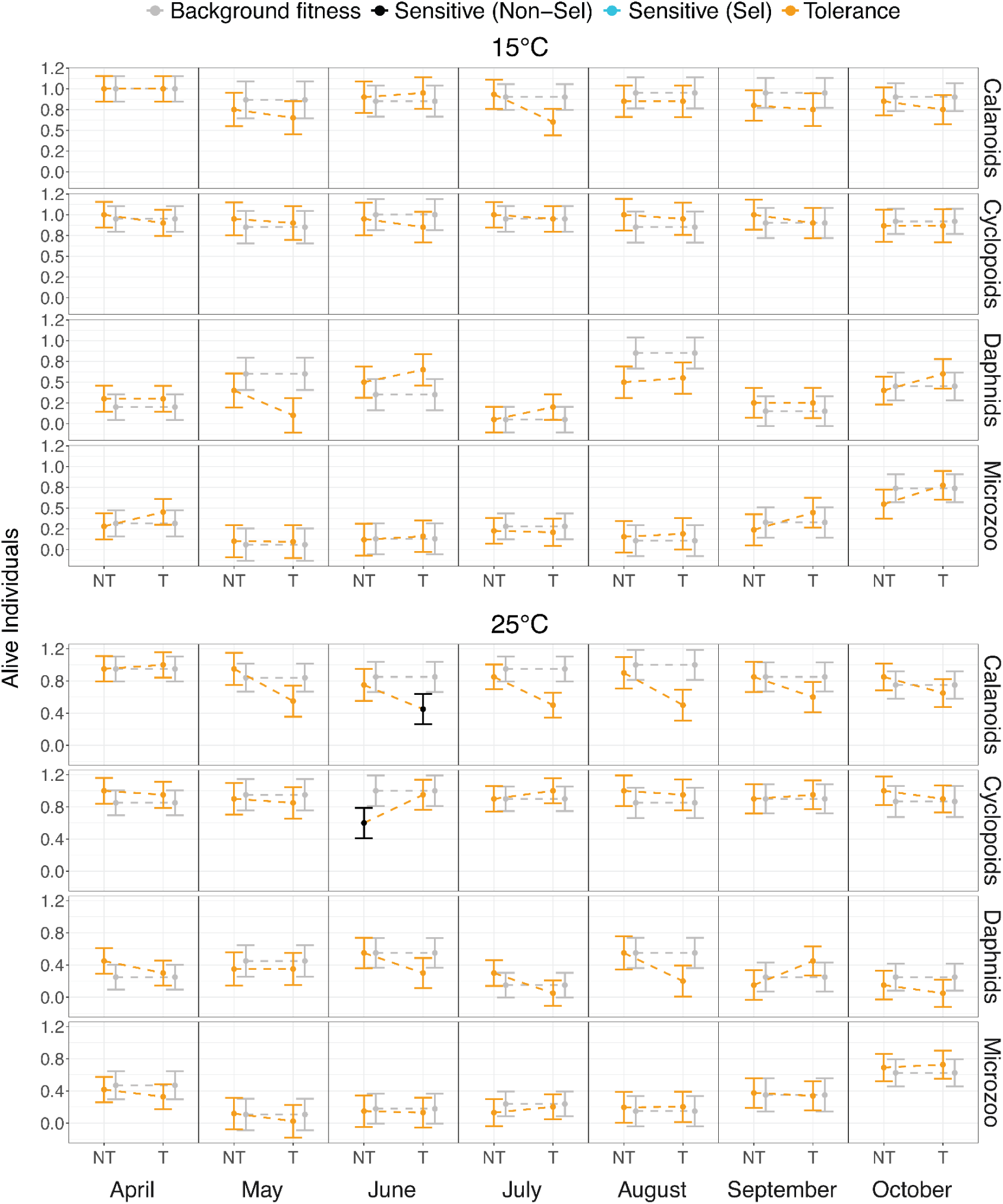
Predicted survival (fitness) of zooplankton. Each panel shows a zooplankton group (rows) over time (columns), with linear-model predictions shown for each combination of food type and temperature. **Grey dots** indicate the mean of control treatments (zooplankton fed only on green algae), while **colored dots** represent the mean of treatments with zooplankton grazing under different food conditions: non-toxic (NT) and toxic (T) cyanobacteria. The color of the dots correspond to the expected fitness outcomes under different zooplankton grazing behaviors (**see** Fig. 2 **B**); specifically black and blue colors denote significant three-way interactions (*zooplankton × food × temperature*) identified through post-hoc testing. *Sensitive (Sel)* indicates selective feeders avoiding cyanobacteria, while *Sensitive (Non-Sel)* indicates non-selective feeders. Error bars indicate 95% confidence intervals of predicted means. All significant post-hoc tests are reported in **Table 2**.

**Table 2:** Results of significant post-hoc tests for the fitness analysis (survival). Comparisons are organised by month (separate models) and treatment interaction. P-values are reported for each significant interaction, with values below 0.05 considered statistically significant.

| Contrast | Zoo | Temperature | Estimate | SE | Df | t.ratio | p.value |
| --- | --- | --- | --- | --- | --- | --- | --- |
| April |  |  |  |  |  |  |  |
| ***Zoo*** |  |  |  |  |  |  |  |
| Calanoids - Daphnids |  |  | 0.683 | 0.045 | 71 | 15.064 | <0.001 |
| Calanoids - Microzoo |  |  | 0.606 | 0.046 | 71 | 13.198 | <0.001 |
| Cyclopoids - Daphnids |  |  | 0.643 | 0.046 | 71 | 13.954 | <0.001 |
| Cyclopoids - Microzoo |  |  | 0.565 | 0.045 | 71 | 12.449 | <0.001 |
| May |  |  |  |  |  |  |  |
| ***Zoo*** |  |  |  |  |  |  |  |
| Calanoids - Daphnids |  |  | 0.403 | 0.058 | 71 | 6.955 | <0.001 |
| Calanoids - Microzoo |  |  | 0.757 | 0.074 | 71 | 10.163 | <0.001 |
| Cyclopoids - Daphnids |  |  | 0.485 | 0.063 | 71 | 7.706 | <0.001 |
| Cyclopoids - Microzoo |  |  | 0.838 | 0.059 | 71 | 14.188 | <0.001 |
| Daphnids - Microzoo |  |  | 0.354 | 0.070 | 71 | 5.041 | <0.001 |
| June |  |  |  |  |  |  |  |
| ***Zoo*** |  |  |  |  |  |  |  |
| Calanoids - Daphnids |  |  | 0.316 | 0.055 | 71 | 5.773 | <0.001 |
| Calanoids - Microzoo |  |  | 0.669 | 0.056 | 71 | 11.983 | <0.001 |
| Cyclopoids - Daphnids |  |  | 0.413 | 0.055 | 71 | 7.561 | <0.001 |
| Cyclopoids - Microzoo |  |  | 0.766 | 0.056 | 71 | 13.800 | <0.001 |
| Daphnids - Microzoo |  |  | 0.353 | 0.055 | 71 | 6.412 | <0.001 |
| **Zoo:Food:Temperature** |  |  |  |  |  |  |  |
| Control - T | Calanoids | 25 | 1.484 | 0.599 | 71 | 2.477 | 0.041 |
| Control - NT | Cyclopoids | 25 | 1.533 | 0.625 | 71 | 2.453 | 0.043 |
| NT - T | Cyclopoids | 25 | -0.370 | 0.134 | 71 | -2.759 | 0.020 |
| July |  |  |  |  |  |  |  |
| **Zoo** |  |  |  |  |  |  |  |
| Calanoids - Cyclopoids |  |  | -0.175 | 0.046 | 70 | -3.770 | 0.002 |
| Calanoids - Daphnids |  |  | 0.638 | 0.047 | 70 | 13.627 | <0.001 |
| Calanoids - Microzoo |  |  | 0.568 | 0.046 | 70 | 12.410 | <0.001 |
| Cyclopoids - Daphnids |  |  | 0.812 | 0.045 | 70 | 18.004 | <0.001 |
| Cyclopoids - Microzoo |  |  | 0.743 | 0.045 | 70 | 16.428 | <0.001 |
| **Zoo:Food** |  |  |  |  |  |  |  |
| Control - T | Calanoids |  | 1.165 | 0.458 | 70 | 2.543 | 0.035 |
| August |  |  |  |  |  |  |  |
| **Zoo** |  |  |  |  |  |  |  |
| Calanoids - Daphnids |  |  | 0.288 | 0.056 | 71 | 5.130 | <0.001 |
| Calanoids - Microzoo |  |  | 0.677 | 0.055 | 71 | 12.322 | <0.001 |
| Cyclopoids - Daphnids |  |  | 0.369 | 0.058 | 71 | 6.364 | <0.001 |
| Cyclopoids - Microzoo |  |  | 0.758 | 0.055 | 71 | 13.727 | <0.001 |
| Daphnids -<br>Microzoo |  |  | 0.389 | 0.056 | 71 | 6.885 | <0.001 |
| September |  |  |  |  |  |  |  |
| **Zoo** |  |  |  |  |  |  |  |
| Calanoids -<br>Daphnids |  |  | 0.550 | 0.053 | 70 | 10.364 | <0.001 |
| Calanoids -<br>Microzoo |  |  | 0.454 | 0.054 | 70 | 8.449 | <0.001 |
| Cyclopoids -<br>Daphnids |  |  | 0.675 | 0.053 | 70 | 12.652 | <0.001 |
| Cyclopoids -<br>Microzoo |  |  | 0.579 | 0.054 | 70 | 10.700 | <0.001 |
| October |  |  |  |  |  |  |  |
| **Zoo** |  |  |  |  |  |  |  |
| Calanoids -<br>Daphnids |  |  | 0.466 | 0.050 | 69 | 9.352 | <0.001 |
| Cyclopoids -<br>Daphnids |  |  | 0.571 | 0.052 | 69 | 10.943 | <0.001 |
| Cyclopoids -<br>Microzoo |  |  | 0.236 | 0.052 | 69 | 4.521 | <0.001 |
| Daphnids -<br>Microzoo |  |  | -0.335 | 0.054 | 69 | -6.195 | <0.001 |

Across all months and treatments, copepods demonstrated consistently higher survival rates compared to daphnids and microzooplankton, particularly evident in April, May, June, July, August, and September (**Fig. 4**). This taxonomic difference in survival suggests that copepods may possess superior physiological or behavioural adaptations for coping with challenging food conditions, including the presence of toxic cyanobacteria.

## 4. Discussion

### 4.1 Non-selective grazing dominates zooplankton-cyanobacteria interactions across the seasonal succession

Our repeated experiments reveal that zooplankton communities in Lake Greifen exhibit predominantly non-selective grazing behaviour toward cyanobacteria throughout the seasonal succession, fundamentally challenging the hypothesis that toxic cyanobacteria are avoided by herbivore grazers. Across broad taxonomic groups—cladocerans, calanoid and cyclopoid copepods, and the diverse microzooplankton—we observed consistent consumption of *Microcystis* strains producing an array of secondary metabolites, including microcystins (MCs) (**Fig. S5**), without a behaviour of systematic avoidance, expected based on previous work (Ger et al., 2016). This generalist feeding pattern persisted despite substantial temporal variation in zooplankton populations (**Fig. S1**), cyanobacterial abundance (peaking in April and October) and secondary metabolite concentrations (anabaenopeptins peaking in August, microcystins in September) (**Fig. 1A**).

The robustness of non-selective grazing is particularly striking when considered against the background dynamics of cyanobacterial secondary metabolites. During periods of elevated cyanotoxin concentrations in the lake, zooplankton maintained feeding rates comparable to control treatments (**Fig. 3**, **Fig. S4**), indicating that the presence of microcystins does not trigger avoidance responses in our case study (note, however, one exception in July under the toxic treatment - **Fig. 3**). Furthermore, the general absence of fitness costs associated with toxic cyanobacteria consumption (**Fig. 4**) suggests that these communities have adapted to enable effective utilisation of cyanobacterial biomass without compromising survival.

### 4.2 Temperature may modulate feeding behaviour through metabolic constraints

Temperature effects on feeding behaviour were generally subtle but revealed important patterns consistent with metabolic hypotheses (Enquist et al., 2015). Contrary to our initial expectations—that higher temperatures would enhance selective preference for green algae (more nutritious) against cyanobacteria (poor food) due to the increased metabolic demands and higher toxicity of MC-producing *Microcystis* (Davis et al., 2009)—all taxonomic groups primarily exhibited non-selective feeding at both temperatures (**Fig. 3**). The persistence of non-selective feeding at higher temperatures (25°C) suggests that increased energetic demands dominate regardless of potential feeding preferences, which could be explained by adaptation to the less nutritious food over time or by the potential costs involved in particle rejection (Olesen et al., 2022; Schultz and Kiørboe, 2009). Interestingly, we also observed two unexpected behavioural shifts at the higher temperature: copepods switched from non-selective to selective consumption of MC-producing cyanobacteria (**Fig. 3**). This suggests that thermal stress may interact with food quality to shape grazing behaviour, and that preferences for food quality may be species-specific (Ger et al., 2010).

In addition to changes in feeding behaviour, another significant temperature-related effect emerged as a fitness cost. In June, both copepod groups exhibited reduced survival under elevated temperature conditions, though in different treatments: calanoid copepods with MC-producing cyanobacteria, and cyclopoid copepods with strains that did not produce MCs, but still had an array of secondary metabolites (**Fig. S5**). This temporal specificity is particularly intriguing, as June marks a transitional period when cyanobacterial densities remain low while toxin concentrations begin to rise (**Fig. 1A**). The restriction of these fitness costs to mainly high temperatures suggests that thermal stress may amplify the effects of bioactive compounds (Claska, 2002; Xiang et al., 2017), whether known toxins like microcystins or potentially unknown secondary metabolites such as non-microcystin cyanopeptides, which are commonly found in Lake Greifen and the used cyanobacterial strains (**Fig. 1A** and **Fig. S5**).

### 4.3 Taxonomic variation in behaviour reveals group-specific patterns

Despite the overall pattern of non-selective grazing, distinct zooplankton taxa exhibited subtle yet statistically significant differences in their responses to cyanobacterial food sources. Daphnids — represented by *Daphnia galeata*, *D. pulicaria*, *D. cucullata*, *D. hyalina*, and their hybrids (the *D. galeata–D. cucullata–D. hyalina* complex) — demonstrated the most consistent non-selective feeding behaviour, consistent with their obligate filter-feeding strategy. However, during periods of peak cyanobacterial abundance (April and September), daphnids unexpectedly exhibited a statistically significant selective preference for non-toxic cyanobacteria over green algae (**Fig. 3**). This suggests that even generalist filter feeders can adjust their feeding behaviour under certain environmental conditions. While surprising given the general assumption of non-selectivity in daphnids, these findings align with previous studies documenting selective feeding in other *Daphnia* species, such as *D. ambigua* (Tillmanns et al., 2011) and *D. pulex* (Meise et al., 1985). Although this may be the first documented case of *Daphnia* from natural populations preferentially selecting non-toxic cyanobacteria, it may not be entirely unexpected. Adapted *Daphnia* clones are commonly found in systems where cyanobacteria are prevalent (Isanta-Navarro et al., 2021; Sarnelle and Wilson, 2005; Schaffner et al., 2019), and previous research has shown that prior exposure to cyanobacteria can enable *Daphnia* to maintain growth when feeding on toxic strains and even reduce cyanobacterial biomass (Chislock et al., 2013; Drugă et al., 2016).

Copepods displayed more complex temporal patterns, contradicting established cyanobacteria avoidance paradigms (Ger et al., 2016). Both calanoid and cyclopoid copepods not only consumed toxic cyanobacteria without discrimination but occasionally preferred cyanobacterial prey over green algae. Notably, calanoid copepods showed statistically significant selective preference for MC-producing cyanobacteria at elevated temperatures in April, while cyclopoid copepods exhibited similar behavior in September (**Fig. 3**). These findings support previous evidence from the Baltic Sea (Novotny et al., 2023, 6 2021) and challenge the conventional view that copepods systematically avoid toxic cyanobacteria, suggesting that local adaptation may override general feeding preferences in systems with persistent cyanobacterial presence. The behavioural change towards selective preference of toxic cyanobacteria under higher temperatures suggests a possible role of adapted copepods in the top-down control of cyanobacteria blooms under climate warming.

Probably as a consequence of its taxonomic heterogeneity—including rotifers, ciliates, nauplii, and dinoflagellates—the microzooplankton assemblage exhibited a remarkably consistent pattern of non-selective grazing and showed no signs of fitness costs across treatments. Seasonal turnover of organisms (**Fig. S1**) may have masked group-specific selective behaviour, for example, for phytoplankton of different sizes (Eyring et al., 2025a). The sole exception to non-selectivity occurred in April and at elevated temperatures (25°C), where microzooplankton displayed a preference for green algae over non-toxic cyanobacteria (**Fig. 3**). The microzooplankton community in April had a remarkably different composition from the rest of the year, with notably lower densities of ciliates compared to later in the season (**Fig. S1**), when non-selective feeding dominated. This suggests that ciliates may play a key role in maintaining non-discriminatory feeding behaviour within the microzooplankton community (Abdullah Al et al., 2023). The temperature-dependent effect may instead reflect the interaction between thermal stress and metabolic constraints in small-bodied organisms, particularly given that ambient lake temperatures in April (8.6°C) were substantially lower than the experimental conditions. A similar pattern has been observed in the rotifer *Brachionus calyciflorus* (Xiang et al., 2017).

### 4.4 Implications for cyanobacterial bloom dynamics and ecosystem management

Our findings have significant implications for understanding cyanobacterial bloom development and persistence in temperate lakes. Lake Greifen has experienced reduced phosphorus loading since nutrient management initiatives began in the 1990s (Bürgi et al., 2003; Merz et al., 2023), resulting in fewer extensive surface blooms over recent decades despite increasing temperatures due to climate change (Tellenbach et al., 2016), and increasing prevalence of cyanotoxins like MC (Monchamp et al., 2016b). The non-selective grazing behaviour and absence of fitness costs we observed suggest that adapted zooplankton communities could potentially influence cyanobacterial bloom dynamics through top-down control, complementing bottom-up nutrient management strategies.

This perspective contrasts with traditional models that emphasise selective avoidance (especially of copepods) as a mechanism promoting cyanobacterial dominance (Ger et al., 2016). We note that most of our assumptions regarding the behaviour of zooplankton and effects of *Microcystis* come from controlled laboratory experiments and single cell strains characterised for the production of microcystins only. In the field, expectations might deviate significantly from laboratory assumptions. Our results support recent empirical evidence that zooplankton can actively regulate cyanobacterial abundances through sustained grazing pressure, even when toxin-producing species are present (Novotny et al., 2023, 6 2021). The selective preference for cyanobacteria during peak abundance periods further suggests that adapted zooplankton may intensify grazing pressure when bloom conditions develop, providing a negative feedback mechanism that could help constrain bloom expansion (Chislock et al., 2013; Sweeney et al., 2022).

### 4.5 Study limitations and future research directions

Some methodological constraints of our findings suggest important directions for future research. Our experimental design focused on short-term feeding behaviour and survival responses, potentially missing longer-term fitness consequences such as reproductive success or multigenerational adaptation dynamics. Sublethal effects of toxin ingestion may accumulate over extended periods, leading to fitness costs that our experimental timeframe could not detect.

The taxonomic heterogeneity within our mesozooplankton and microzooplankton categories, while reflecting natural community composition, may obscure species-specific responses that could be ecologically important. Seasonal shifts in community composition add another layer of complexity, as the relative abundance of species with different physiological tolerances likely influenced our group-level results. Future studies incorporating species-level resolution and longer experimental durations would provide more mechanistic insights into the adaptive processes we observed. These adaptive processes naturally include physiological plasticity and sorting of different genetic lineages: the role of these potential mechanisms in the patterns we have observed requires a targeted investigation.

Despite these limitations, our findings open several critical research avenues that could reshape our understanding of cyanobacterial bloom dynamics. Long-term studies are needed to assess whether the adaptive responses we observed represent stable evolutionary changes or plastic behavioural or physiological adjustments that may vary across years or bloom intensities. Cross-system comparisons would help determine whether our findings from Lake Greifen—a system with a long history of cyanobacterial presence but reduced bloom frequency — are generalizable to lakes with different management histories and bloom patterns.

The role of the other secondary metabolites produced by our studied cyanobacteria (**Fig. S5**) in mediating grazer-cyanobacteria interactions remains poorly understood and warrants further investigation. Our observation that copepods sometimes experienced reduced survival with non-MC-producing cyanobacterial strains suggests that the dichotomy between “toxic” and “non-toxic” cyanobacteria may oversimplify the complex chemical ecology of these interactions (Janssen, 2019). Future studies should characterise the full metabolite profiles of cyanobacterial strains and assess their individual and synergistic effects on grazer performance and behaviour.

From a management perspective, these results emphasise the importance of focusing on biotic interactions alongside traditional nutrient management approaches. The temporal plasticity in zooplankton behaviour that we observed underscores the dynamic role of biotic interactions in regulating cyanobacterial blooms and highlights the need to incorporate both ecological and evolutionary feedbacks into predictive ecological models (Ntetsika et al., 2025). If adapted zooplankton communities can indeed regulate cyanobacterial populations through sustained grazing pressure, conservation and enhancement of zooplankton diversity and abundance (especially of previously adapted species) may represent an effective tool for bloom prevention or control. This biological control may be particularly valuable under climate change scenarios where warming temperatures are expected to favour cyanobacterial growth while simultaneously increasing zooplankton metabolic demands.

In conclusion, our study provides evidence that zooplankton-cyanobacteria interactions are more complex and dynamic than traditionally perceived, with important implications for understanding and managing freshwater ecosystem stability under global change.

## Funding sources

This research was funded by the Swiss National Science Foundation (project 202290 Cyanobloom), with the contribution for infrastructure by the Swiss Federal Office for the Environment (contract Nr. Q392-1149).

## CRediT authorship contribution statement

**Pinelopi Ntetsika:** Conceptualization, Data curation, Formal analysis, Investigation, Methodology, Project administration, Supervision, Validation, Visualization, Writing – original draft, Writing – review & editing. **Salomé Stauffer:** Data curation, Formal analysis, Investigation, Writing – first draft, Writing – review & editing. **Stefanie Eyring:** Resources, Writing – original draft, Writing – review & editing. **Marta Reyes:** Resources, Writing – original draft, Writing – review & editing. **Xuejian Wang:** Resources, Writing – original draft, Writing – review & editing. **Elisabeth M.-L. Janssen:** Resources, Writing – original draft, Writing – review & editing. **Francesco Pomati:** Conceptualization, Formal analysis, Funding acquisition, Investigation, Methodology, Project administration, Supervision, Validation, Writing – original draft, Writing – review & editing.

## Declaration of competing interest

The authors declare that they have no known competing financial interests or personal relationships that could have appeared to influence the work reported in this paper.

## Acknowledgements

We are grateful to **Silvana Käser** for her valuable assistance in setting up and operating the equipment used in this study, and for dedicating considerable time in explaining how to accurately distinguish zooplankton species. We thank **Prof. Dr. Jukka Jokela** for his invaluable support in data analysis and for providing critical feedback on both experimental design and manuscript revision.

## Data availability

All codes and a subset of the dataset are currently available publicly on https://github.com/PinelopiNt/ZooFeedingBehaviour. Upon request, we can provide access to the full dataset. All experimental data and statistical code will be deposited on https://opendata.eawag.ch/ upon acceptance, in accordance with the journal’s data sharing policies.

## Notes

### Competing Interest Statement

The authors have declared no competing interest.

